## Supplementary materials for "Gender-based disparities and biases in science: an observational study of a virtual conference"

### Supplementary Note

May 5, 2022

#### Contents

**1 Supplementary Figures** **2**

**2 Supplementary Tables** **4**

**3 Observation Guidelines and Form** **7**

**4 Interview Guide** **13**

**5 Interview Consent Form** **17**

### 1 Supplementary Figures

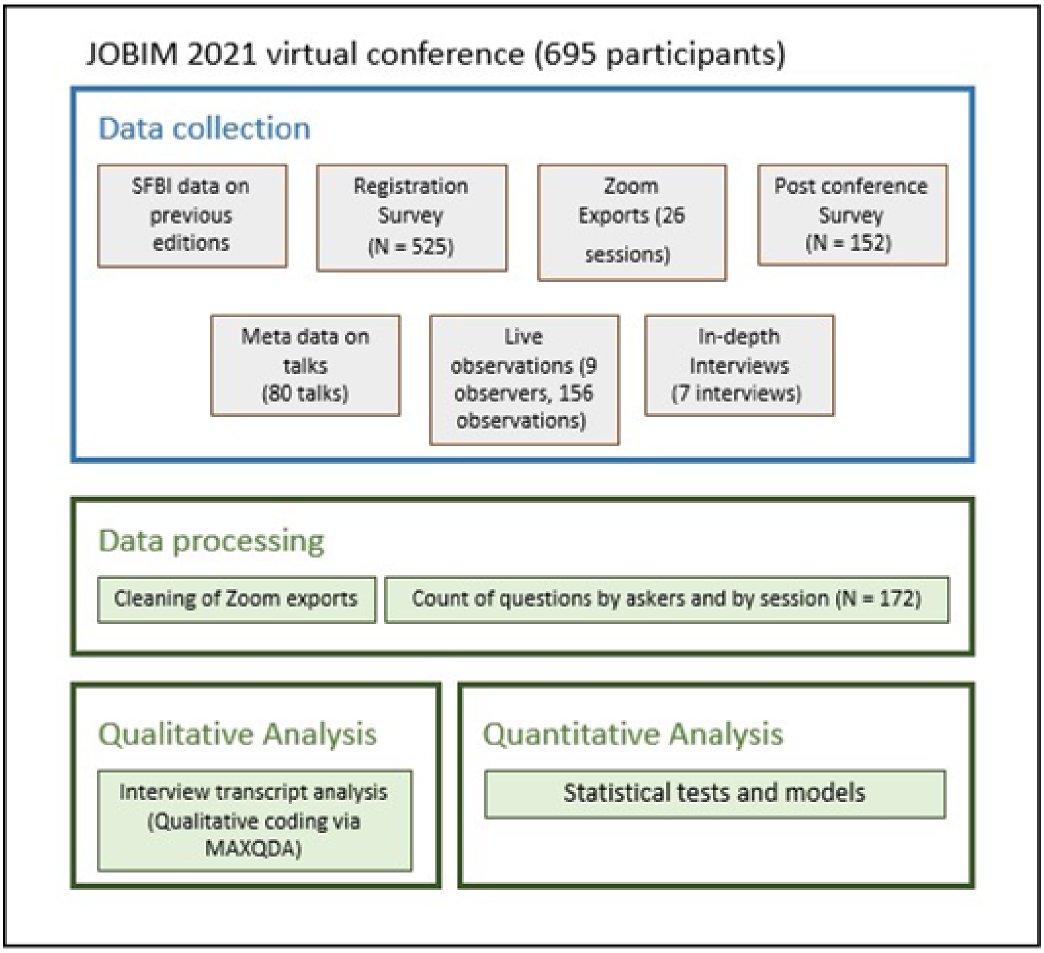

**Figure S1**: **Overview of the study** with the list of data collected, the main data processing steps and the main analysis steps.

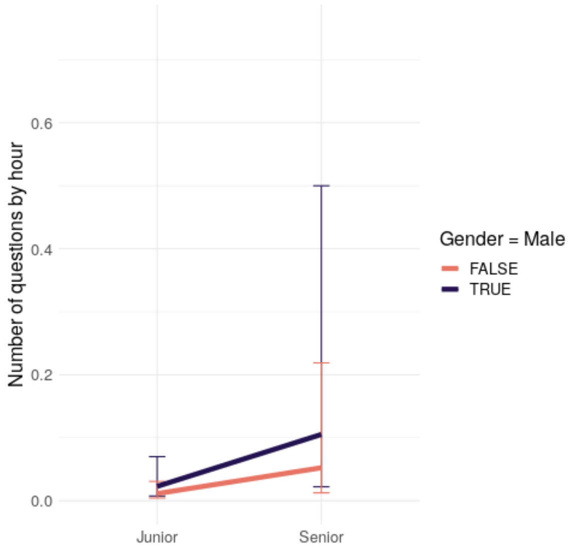

**Figure S2**: **Predicted Mean rate for Junior and Senior academics conditioned by their gender**. By senior, we refer to an attendee older than 35 and with a permanent position. By junior, we refer to an attendee younger than 35 and with a short term contract.

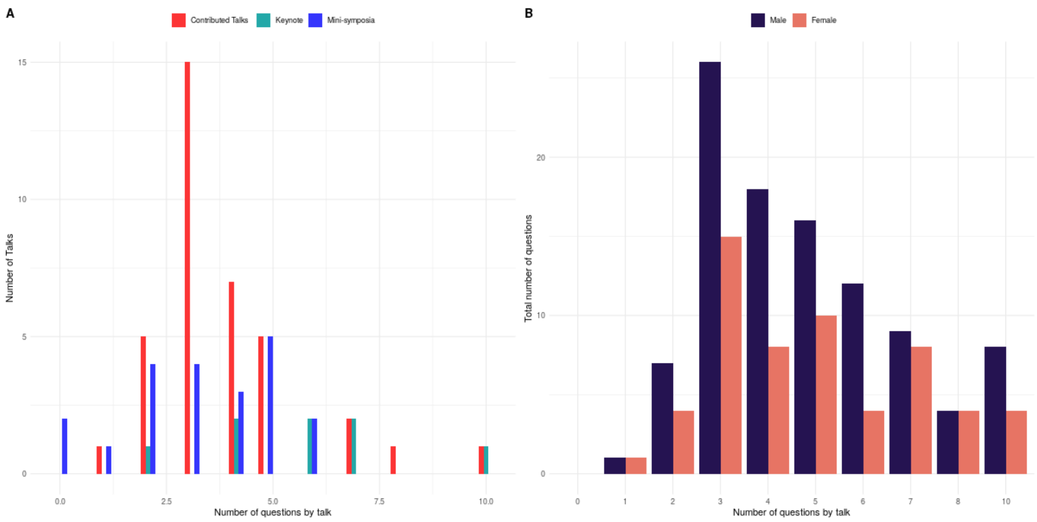

**Figure S3**: **Effect of the length of the question session** A) Histogram of the number of questions asked at the end of the talk by type of sessions (color coded), B) Total number of questions asked by gender (indicated by the color of the bars) with respect to the total number of questions asked at the end of the talk (used as a proxy for the duration of question session).

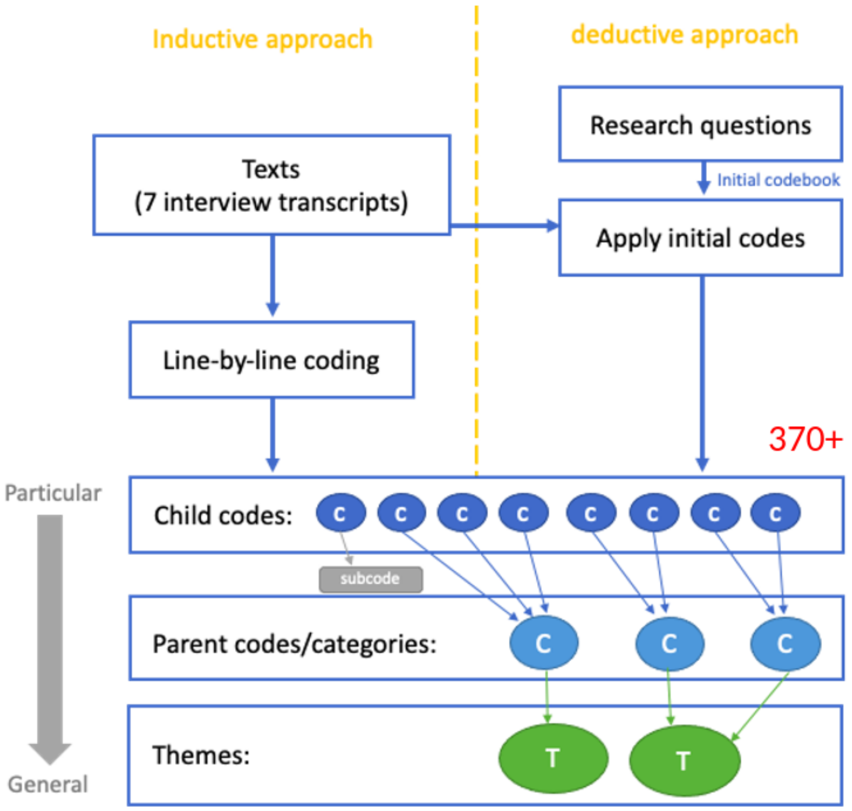

**Figure S4**: **Qualitative Analysis of interview transcripts**. Each line of the transcript is assigned to a child code either derived from the research questions or created to represent the recurring topics. When all the transcripts have been processed, the resulting child codes are gathered into broader categories : parent codes. Finally parent codes are summarized in themes.

### 2 Supplementary Tables

| | Service Platforms | Research | Networking and Working group | Total |
|---|---|---|---|---|
| Female | 10 | 57 | 3 | 69 |
| Male | 11 | 55 | 3 | 70 |

**Table S1**: **Number of poster by gender and poster categories**

| Model | df | AIC |
|---|---|---|
| $\ln(\mathrm{Y}) = \beta_0 + \beta_1 \times (Gender = Male) + \varepsilon$ | 2 | 660.44 |
| $\ln(\mathrm{Y}) = \beta_0 + \beta_1 \times (Age \geq 35)$<br>$+ \beta_2 \times (ProfessionalStatus = permanent) + \varepsilon$ | 3 | 623.63 |
| $\ln(\mathrm{Y}) = \beta_0 + \beta_1 \times (Gender = Male)$<br>$+ \beta_2 \times (Age \geq 35) + \beta_3 \times (ProfessionalStatus = permanent) + \varepsilon$ | 3 | 615.69 |
| $\ln(\mathrm{Y}) = \beta_0 + \beta_1 \times (Gender = Male)$<br>$+\beta_2 \times (Age \geq 35) + \beta_3 \times (ProfessionalStatus = permanent) + \varepsilon$ | 4 | 610.46 |
| $\ln(\mathrm{Y}) = \beta_0 + \beta_1 \times (Gender = Male) + \beta_2 \times (Age \geq 35)$<br>$+ \beta_3 \times (ProfessionalStatus = permanent) + \beta_4 \times (Age \geq 35) \times (Gender = Male) + \varepsilon$ | 5 | 611.66 |
| $\ln(\mathrm{Y}) = \beta_0 + \beta_1 \times (Gender = Male) + \beta_2 \times (Age \geq 35)$<br>$+ \beta_3 \times (ProfessionalStatus = permanent)$<br>$+ \beta_4 \times (ProfessionalStatus = permanent) \times (Gender = Male) + \varepsilon$ | 5 | 609.9 |

**Table S2**: **AIC by variables included in the Poisson regression**

| Child Codes | Categories/Parent Codes | Themes |
|---|---|---|
| Discouraged to ask questions | Discouragement | Women and gender minority report negative experiences based on sexual orientations, gender identities and expressions |
| Discouraged to choose a scientific career | | |
| Labelling | Professional ability being undermined | |
| Alienation | | |
| Gender identity impacts confidence | Gender identity impacts confidence | Correlation between gender identity, confidence and career advancement |
| Confidence impacts career advancement | Confidence impacts career advancement | |
| Confidence improves through self-acceptance | How confidence is improved | |
| Confidence improves through experiences | | |
| Conscious about gender issues | Attitude: conscious | Women and gender minorities are more proactive to challenge gender inequalities in the workplace |
| Conscious about micro-aggression | | |
| Conscious about gender privilege | | |
| Curious to learn more | Attitude: curious | |
| Self-identify as an ally | Attitude: supportive | |
| Self-identify as a feminist | | |
| Eager to make a change | Attitude: active | |
| Action-taking | | |
| Unaware of micro-aggression | Attitude: oblivious | |
| Men do not feel concerned | | |
| Men lack of actions | | |
| Personal experience | Objects of discrimination and harassment | Women are subjected to gender-based discrimination and sexual harassment |
| Happened to colleagues | | |
| Sexual harassment towards women | Types of discrimination and harassment | |
| Gender-based discrimination towards women | | |
| Intersectional discrimination | | |
| Leaky pipeline | Typical discriminative phenomena are identified | Other findings |
| Childcare burden on women / motherhood discrimination | | |
| Male-dominant environment | | |
| Glass ceiling | | |
| "Positive" discrimination regarding quota | | |
| Positive changes toward gender equality | Positive changes are observed | |
| Field differences in gender imbalance | Field differences in gender imbalance | |
| Active in social events | Activeness in conference participation | |
| Active in poster sessions | | |
| Lack of support system | Institutional efforts on gender equality | |
| No channel to speak up | | |
| Counselling sessions for students only | | |
| Gender equality report | | |
| Lack of positions | Self-identified career obstacles | |
| Lack of self-confidence | | |
| Lack of fundings | | |
| Negative: hard to build long-term connections | Past experiences with conferences | |
| Negative: lack of useful information | | |
| Negative: difficult when working in industry | | |
| Negative: not a fan of big conferences | | |
| Positive: enjoy hanging out with colleagues | | |
| Positive: networking opportunities | | |
| Positive: opportunity to ask questions | | |
| Positive: boost new ideas | | |
| Networking and/or collaboration | Motivation for attending conferences | |
| Self-improvement | | |
| Socialisation | | |
| Re-connect with previous colleagues | | |
| getting up to date with the field | | |
| Self-advertisement/visibility | | |
| Biology | Professional field | |
| Bioinformatics | | |
| Structural biology | | |
| Computer science | | |
| 6 years | Years in STEM | |
| 7-8 years | | |
| 10-12 years | | |
| 10+ years | | |
| 16 years | | |
| 25 years | | |

**Table S3: Final Codebook**

| Child Codes | Categories/Parent Codes |
|---|---|
| Professional field of work | Demographic information |
| Years in STEM | |
| Conference attendance frequency | Conference/Question-asking |
| Number of times attending JOBIM | |
| View on the importance of conferences | |
| Motivations for attending conferences | |
| Activeness in conferences | |
| In-person vs virtual conferences | |
| Gender balance/imbalance in STEM | Gender |
| Discrimination/Harassment | |
| Diversity and inclusion in STEM | Other |
| Self-identified career obstacles | |

**Table S4: Initial Codebook**

| identified by first name / Self identified | Female | Male | Not identified |
|---|---|---|---|
| Agender | 1 | 0 | 0 |
| Female | 223 | 1 | 34 |
| Male | 2 | 217 | 29 |
| Non-binary | 2 | 2 | 0 |

**Table S5**: **Confusion Matrix between gender identified from first name versus self identified gender.** Each case is the count of attendees belonging to this category

### 3 Observation Guidelines and Form

**Planning for observations during the JOBIM 2021 Conference**

JOBIM 2021 pilot project – "Gender Speaking Differences in Academia"

*(for internal use only and used by the observers of this study and conference moderators)*

#### I. Observational guidelines – instructions for collecting observational data

**Background**: This observational study is designed as part of the pilot project that focuses on gender speaking differences in academic. Since some studies have shown significant disparities between women and men in question-asking behaviours in academic seminar, we propose to perform the evaluation on gender speaking differences during the JOBIM 2021 Conference. Through this study, we hope to further discover the factors that contribute to gender-based biases in scientific conferences as well as the potential solutions for alleviating such inequality. Since the JOBIM 2021 Conference takes place virtually, the study will provide us with extra insights on disparities between men and women in question-asking behaviours in online settings. Our preliminary research on the existing literature has shown findings including female participants asking fewer questions than male participants in academic seminars. Through in-depth discussions with seminar participants and organisers at the Institut Pasteur, we acknowledge that such a phenomenon was commonly present in previous scientific seminars/conferences.

*The design of this planning is based on a set of hypotheses and research questions (attached in the Appendix[1])

**Approach:** In order to confirm the gender disparity in large scientific conferences and to delve deeper into the factors that possibly contribute to the unequal participation, we adopt an evident-based and ethnographic approach to this study. Participant observation is the key access to first-hand data. The following instructions for observations are designed to facilitate the observers' note-taking during each session and to record the data collection process.

**Remark:** The planning for observations is for internal use only and used by observers of this study and conference moderators. We recommended the viewers of this document to avoid sharing the detailed information with your colleagues or other conference attendees. This is to reduce the impact of this observation in the behaviours of the observees. We reply on your understanding and assistance to ensure the quality of our research.

#### II. Instructions for observations

The instructions are used by the observers of the study and conference moderators. The instructions for observing a session (keynote session/paradelle session) provide observational guideline for the following stages – 1) prior to the session, 2) during the session, and 3) after the session.

*Pilot testing of the observational sheet was performed. The testing confirmed that **at least 2 observers** are needed to observe one session simultaneously.

[1] Page 6

- Prior to the observation session:
  - Please make sure that you are familiar with the data collection procedure.
  - If possible, fill in some parts of the observational sheet (see the following section) based on information that you already have. For example: date, name of the observer, type of the session, name of the chairperson, name of the moderator(s), etc.

- During the session:
  - To observers: please fill in the following observational sheet basing on your observation. All fields are mandatory.
  - To moderators: please fill in the fields in blue in the following observational sheet, or provide the information in other ways (e.g., in a word document, or via emails).

We separate questions into two groups – interrupting questions (questions asked during the talk) and questions asked after the talk, although we acknowledge that for most of the sessions the attendees will not be able to interrupt the talks. Please be aware that a question can be a comment, a statement, or a statement that particularly solicits a response. Definitions of items in the observational sheet are listed as follows:

**Date**: the date of the session as DD/MM/YY, e.g., 09/07/21.
**Time**: the starting time of the talk as HHMM, e.g., 1430.
**Observer**: the name of the observer (the name will be anonymized).
**Type of session**: the typical types of sessions in the JOBOM Conference are keynote sessions and parallel sessions. If the observed session is neither of the two, please specify.
**Title of session**: the tile of the session (the title will be anonymized).
**Speaker**: the name of the speaker (the name will be anonymized).
**Chairperson**: the name of the chairperson (the title will be anonymized).
**Moderator 1**: the name of the first moderator (the name will be anonymized).
**Moderator 2**: the name of the second moderator (the name will be anonymized).
**N_attendees**: the number of attendees at the session.
**N_male**: the number of male attendees at the session.
**N_female**: the number of female attendees at the session.
**Asker**: the name of the attendee who asks the question (the name will be anonymized).
**Gender**: the observed gender of the person – male, female or N/A (if the information is not clear/available).
**Question_time_start**: the time at which the question starts as HHMM, e.g., 15:45.
**Question_time_end**: the time at which the question ends as HHMM, e.g., 15:46.
**Question type:** the question should be categorized as 1) Challenges, 2) Clarification, 3) Information-seeking, 4) Receptions, 5) Compliments / Containing compliments, or 6) Others______ / Not sure. Please use to the number to specify the type of the questions. One question can be classified into one or more of the above-mentioned categories.
**Remarks**: Please note down anything else that could be relevant to this study or provide complementary information.
**N_questions**: the total number of questions asked during the session.
**N_male_asker**: the total number of male askers.
**N_female_asker**: the total number of female askers.

#### 1. Basic information of the session[2]

Date: ______________________ Time[3]: ____________________
Observer: ________________

Type of session: Keynote ☐ Parallel ☐ Other__________

Title of session: ________________________________________

Speaker: __________________ male ☐ / female ☐ / NA ☐

Chairperson: _________________ male ☐ / female ☐ / NA ☐

Moderator 1: _________________ male ☐ / female ☐ / NA ☐

Moderator 2: _________________ male ☐ / female ☐ / NA ☐

#### 2. Attendees of the session

N_attendees: ☐ N_male: ☐ N_female: ☐

* Please calculate the number of attendees 15 minutes after the session has started

Name list of the attendees:

* Please extract the name list 15 minutes after the session has started

#### 3. Question-answer behaviours

- Interrupting questions (during the talk)

| Number | Asker | Gender | Question_ time_start | Question_ time_end | Question_ type[4] | Remarks |
|---|---|---|---|---|---|---|
| Question_1 | | male☐ / female☐ / NA☐ | | | | |
| Question_2 | | male☐ / female☐ / NA☐ | | | | |

[2] All identifications of gender in this observational sheet refer to “observed gender”.
[3] Starting time of the observation.
[4] Categorizations: 1) Challenges; 2) Clarification; 3) Information-seeking; 4) Receptions; 5) Compliments / Containing compliments; 6) Others______ / Not sure

3

- Questions asked after the talk

| Number | Asker | Gender | Question_ time_start | Question_ time_end | Question_ type | Remarks |
|---|---|---|---|---|---|---|
| Question_1 | | male☐ / female☐ / NA☐ | | | | |
| Question_2 | | male☐ / female☐ / NA☐ | | | | |
| Question_3 | | male☐ / female☐ / NA☐ | | | | |
| Question_4 | | male☐ / female☐ / NA☐ | | | | |

- Total questions asked

N_questions: ☐ N_male_asker: ☐ N_female_asker: ☐

**4. Questions in the chat**

- Messages in the public chat box with details, including times and names (copy all messages from the chat box in the end of the session):

- Messages sent privately to the moderators with details, including times and names:

- <u>After the session:</u>
  - Please make sure the competed observational sheets and collected data are stored in a safe place.
  - Send all the information to the JOBIM pilot study team once you finish your tasks as an observer/moderator.

4

#### III. Contact information

For questions and more information, please contact contact us at. To get in touch with individual researchers of the project, please refer to the following contact list:

Junhanlu Zhang
Eng. Rachel Torchet
Dr. Hanna Julienne

An observational training session will soon be organized for observers.
A Q&A session may be organized for moderators too, depending on the requests we receive. We will keep you updated once such sessions are planned.

#### *Appendix*

| Hypotheses/research questions | Observational data | Remarks |
|---|---|---|
| 1. Do men ask more questions than other genders during keynote and poster sessions? | • Numbers of question asked | |
| | • Observed gender of the asker (male/female/NA) | |
| | • Name of the asker (if not anonymous) | For later use to compare the observed gender with the self-identified gender. |
| 2. What is the gender ratio of the session and is it a factor that influence participants' question-asking behaviours? | • Total number of participants | |
| | • Number of participants by their observed genders (male/female/NA) | |
| | • Name list of the participants | For later use to compare the observed gender with the self-identified gender. |
| 3. Are interrupting questions[5] more likely to be asked by men? | • Observed gender of the person who asks interrupting questions | |
| | • Name of the person who asks interrupting questions | For later use to compare the observed gender with the self-identified gender. |
| 4. Are the first questions[6] more likely to be asked by men? | • Observed gender of the person who asks the first question | |
| | • Name of the person who asks the first question | For later use to compare the observed gender with the self-identified gender. |
| 5. Do men take more time to ask questions than women and other genders do? | • Starting time of the question asked | |
| | • Ending time of the question asked | |
| | • Observed gender of the asker | |
| | • Name of the asker | For later use to compare the observed gender with the self-identified gender. |
| 6. Do the genders of the speaker and moderator impact question-asking behaviours of the participants? | • Observed gender of the speaker | |
| | • Name of the speaker | For later use to compare the observed gender with the self-identified gender. |
| | • Observed gender of the moderator | |
| | • Name of the moderator | For later use to compare the observed gender with the self-identified gender. |
| 7. Are there emotional differences in question-asking behaviours based on the gender of the asker? | • Categorization of each question asked | - Challenges<br>- Clarification<br>- Information-seeking<br>- Receptions<br>- Compliments / Containing compliments<br>- Others / Not sure |

[5] Questions that interrupt the speaker during his/her talk.
[6] First questions asked after a talk or presentation.

### 4 Interview Guide

#### Interview guide (with interview questions)

**Research project title**
JOBIM 2021 Pilot Project - Gender Speaking Differences in Academia

**Research investigators**
Junhanlu Zhang, Rachel Torchet & Hanna Julienne

---

This interview guide is developed to assist the interviews as part of the JOBIM 2021 Pilot Project. The guide consists of guidelines throughout three stages of the interview procedure – preparation, interview, and post-interview, as well as interview questions covering three major areas – demographic information, experience with academic conferences, and gender-related questions in STEM. Basic information of the interviews is as following,

**Type of interviews:** semi-structured in-depth interviews
**Interviewees:** attendees of the JOBIM 2021 conference who agreed at the conference registration stage to contribute to this study
**Interviewee sample size:** 5-7 interviewees
**Time of interviews:** 30-45 minutes
**Interview channel:** videoconferencing platforms (e.g., Teams and Zoom)

---

##### Stage 1: Preparation

- Invite potential interviewees to join the interviews and schedule the time/date once the interview is confirmed.
- Send an interview reminder to the interviewee with the consent form for both parties to sign 1-3 days prior to the agreed interview time.
- If applicable, pay attention to the interviewee's registration and post-survey data to adapt the interview accordingly.
- Prepare the equipment (e.g., notebook and pen) to take complimentary notes and to make backup recording (e.g., recording pen or phone) during the interview.

##### Stage 2: Interview

<table>
<tr><td>Introduction</td><td colspan="2">- Introduce the project and the purpose of the interview.<br><br>- Emphasize on the confidentiality of the interview, and double check with the interview if they understand the information provided on the consent form.<br><br>- Let the interviewee know that they can interrupt to ask questions anytime during the interview. If there is any question that makes the interviewee uncomfortable, they can refuse to answer the question, or even stop the interview if necessary.<br><br>- Asked for permission to record and start recording.</td></tr>
<tr><td rowspan="2">Research questions</td><td>Demographic information</td><td>- Hong long have you been studying and/or working in the STEM fields?<br><br>- Which is your specialised area of work?<br><br>- What is your current institution?<br><br>- How settled are you in your current position?<br><br>- How is it like working in STEM in general?</td></tr>
<tr><td>Experience with academic conferences</td><td>- How often do you participate in academic conferences either in your professional field or in other fields?<br><br>- Is attending conferences important to you? Why?<br><br>- How do you like such an event based on your past experiences?<br><br>- What usually motivate you to attend a conference?<br><br>- How active are you in academic conferences? For example, do you ask questions?<br><br>- Is it usually easy for you to ask questions during the conference? Why?</td></tr>
</table>

| | | |
|---|---|---|
| | | - Have you participated in online conferences since the pandemic started and how did you like them?<br><br>- Are online and offline conferences different in your opinion? What is your preference?<br><br>- What are your experiences when it comes to meetings and academic seminars? Is it easy to express yourself and how are your ideas received usually? |
| | Gender-related questions in STEM | - What are/were the biggest obstacles that you recognise throughout your career development?<br><br>- Based on your observation, do you see an inequality or imparity between men and women in the STEM fields?<br><br>- The existing literature often considers women as the minority and underprivileged population in STEM, do you agree? Why?<br><br>- What is your general impression on gender diversity and inclusion in STEM?<br><br>- Do you think your gender identity or sexual orientation matters in your work and/or professional life? If you do, in what way does it intervene your work and/or professional life?<br><br>- Have you experienced or observed any type of gender-based discrimination and harassment in conferences or in the workplace? |

| | | |
|---|---|---|
| | | - In your opinion, should we do more to reach gender diversity, equity, and inclusion in STEM? Any ideas or advice? |
| Conclusion | - Ask the interviewee if there is any question or anything to add.<br><br>- Stop the recording.<br><br>- Express appreciation, and let the interviewee know that they can get in touch in the future if there is anything that they would like to address after the interview. Provide contact information if needed. | |

##### Stage 3: Post-interview

- Check notes and add necessary information (e.g., observations during the interview, clarification on notes, or reflections as the interviewer) as soon as possible after the interview is over.
- Check the quality of recording and make sure that the recording is kept in a secure place.

### 5 Interview Consent Form

#### Interview Consent Form

**Research project title**
JOBIM 2021 Pilot Project - Gender Speaking Differences in Academia

**Research investigators**
Junhanlu Zhang, Rachel Torchet & Hanna Julienne

---

Participant's name: ______________________ Interview Date: _____ /_____ /________

---

**Description of the project**

In the JOBIM 2021 conference, we are launching a pilot project on gender speaking differences in academia (https://research.pasteur.fr/en/project/jobim-2021-pilot-project-gender-speaking-differences-in-academia/). Through this evidence-based and mixed-method study, we intend to answer the following question: how to create conditions for gender-equal expression in scientific conferences?

---

Thank you for agreeing to be interviewed as part of the above research project. This interview will take approx. 30-45 minutes. We do not anticipate that there are any risks associated with your participation. However, sensitive personal information may be involved during the interview, and you have the rights to stop the interview or withdraw from the research at any time.

Based on the GDPR regulations, this consent form is necessary for us to ensure that you understand the purpose of your involvement and that you agree the conditions of your participation. Would you therefore read the accompanying **information sheet** and sign this form to certify that you approve the following:

- The interview will be recorded, and a transcript will be produced.
- You will have the access to the transcript as well as the opportunity to correct errors.
- The transcript of the interview will be analysed by Junhanlu Zhang as research investigator.
- Access to the interview recording and transcript will be limited to research investigators of this project – Junhanlu Zhang, Rachel Torchet and Hanna Julienne.
- The information from your interview will be processed and published anonymously. Care will be taken to ensure that other information in the interview that could identify yourself is not revealed.
- The actual recording will be deleted once the project is completed (by the end of the year 2021).
- Any variation of the conditions above will only occur with your further explicit approval.

By signing this consent form, I agree that:

- I am participating in this research project voluntarily, and I understand that I can stop the interview or withdraw from the project at any time.

- I understand that I can exercise other individual rights in compliance with the GDPR regulations – I can access the transcript, object to the processing of the interview data, and I also have the right to the portability of the interview data.

- I have carefully read the information sheet.

- I do not expect any benefit or payment for my participation.

- I understand that I can express myself freely and ask questions whenever I need to during the interview.

- I understand that I am free to contact research team with any question I may have regarding this project in the future.

-----------------------------------------------------------------------------------------------------------------------

| Participant's signature | Researcher's signature |
|---|---|
| Date signed | Date signed |
| _____/_____/________ | _____/_____/________ |

Institut Pasteur

JOBIM 2021 06-09 JUIL INSTITUT PASTEUR | PARIS
